# DeepForest: A Python package for RGB deep learning tree crown delineation

**DOI:** 10.1101/2020.07.07.191551

**Authors:** Ben. G. Weinstein, Sergio Marconi, Mélaine Aubry-Kientz, Gregoire Vincent, Henry Senyondo, Ethan White

**Affiliations:** Department of Wildlife Ecology and Conservation, University of Florida, Gainesville, Florida, USA; AMAP, IRD, CNRS, INRA, Univ Montpellier, CIRAD, 34000 Montpellier, France

**Keywords:** Remote Sensing, Forests, Tree Crowns, Crown Delineation, NEON, Deep learning, RGB

## Abstract

1. Remote sensing of forested landscapes can transform the speed, scale, and cost of forest research. The delineation of individual trees in remote sensing images is an essential task in forest analysis. Here we introduce a new Python package, DeepForest, that detects individual trees in high resolution RGB imagery using deep learning.
2. While deep learning has proven highly effective in a range of computer vision tasks, it requires large amounts of training data that are typically difficult to obtain in ecological studies. DeepForest overcomes this limitation by including a model pre-trained on over 30 million algorithmically generated crowns from 22 forests and fine-tuned using 10,000 hand-labeled crowns from 6 forests.
3. The package supports the application of this general model to new data, fine tuning the model to new datasets with user labeled crowns, training new models, and evaluating model predictions. This simplifies the process of using and retraining deep learning models for a range of forests, sensors, and spatial resolutions.
4. We illustrate the workflow of DeepForest using data from the National Ecological Observatory Network, a tropical forest in French Guiana, and street trees from Portland, Oregon.

## Introduction

Airborne individual tree delineation is a central task for forest ecology and the management of forested landscapes. The growth in sensor quality and data availability has raised hopes that airborne tree maps can complement traditional ground-based surveys (Hamraz et al. 2016). Most approaches to tree delineation in remote sensing use three-dimensional LIDAR data (Coomes et al. 2017), which is currently available for only a small fraction of the Earth’s surface. In contrast, high resolution RGB data has widespread coverage from commercial and government sources and is readily collected using unmanned aerial vehicles. As a result, there is an increasing need for RGB-based tree delineation approaches with easy to use open-source implementations.

The introduction of deep neural networks has greatly enhanced the performance of remote sensing solutions for detecting objects in geospatial images (Zhu et al. 2017). Deep learning models use a series of hierarchical layers to learn directly from training data instead of using expert designed features. Initial layers learn general representations, such as colors and shapes, and subsequent layers learn specific object representations. There are several barriers to applying deep learning to ecological applications including insufficient technical expertise, a lack of large amounts of training data, and the need for significant computational resources. DeepForest provides easy access to deep learning for tree delineation by creating a simple interface for training object detection models, using them to make predictions, and evaluating the accuracy of those predictions. DeepForest also includes a prebuilt model (based on Weinstein et al. 2020) pre-trained on tens of millions of LiDAR generated crowns and fine-tuned using over 10,000 hand-labeled crowns from diverse forests in the National Ecological Observatory Network. Users can apply this model to detect trees in new imagery or provide additional hand-labeled data to fine-tune performance for a specific site or forest type. Predictions from the model for an average 1km2 tile can be made in 7 minutes on a single CPU and DeepForest has built-in support for running on GPU resources to dramatically increase the speed of prediction at large scales.

## DeepForest Software

DeepForest is an open source (MIT license) Python package supporting Python 3.6 and Python 3.7 and has been tested on Windows, macOS, and Linux operating systems. It can be installed using the Python Package Index (https://pypi.org/project/deepforest/) or using the conda package manager for Windows, Linux and OSX (https://github.com/conda-forge/deepforest-feedstock). The software is openly developed on GitHub (https://github.com/weecology/DeepForest) with automated testing and each release is archived on Zenodo (https://doi.org/10.5281/zenodo.2538143). All DeepForest functions are documented online with reproducible examples (https://deepforest.readthedocs.io/) and video tutorials.

### Prebuilt model

DeepForest currently includes one prebuilt model (available by running `deepforest.use_release`) that was trained on data from the National Ecological Observatory Network (NEON) using a semi-supervised approach outlined in Weinstein et al. (2019, 2020) (Figure 1). The model was pretrained on data from 22 NEON sites using an unsupervised LiDAR based algorithm (Silva et al. 2016) to generate millions of moderate quality annotations for model pretraining. The pretrained model was then retrained based on over 10,000 hand-annotations of RGB imagery from six sites (MLBS, NIWO, OSBS, SJER, TEAK, LENO; see NEON site abbreviations S1). The full workflow is shown in Figure 1. While LIDAR data is used to facilitate data generation for the prebuilt model, prediction relies only on RGB data, allowing the model to be used to detect trees using RGB imagery alone. This prebuilt model extends the methods from Weinstein et al. (2019, 2020) by pretraining on a much larger number of trees (30 million LIDAR-generated crowns compared to 10 million in Weinstein et al. 2020) and diversity of sites (22 instead of 4 in Weinstein et al. 2020). Additional details on the modeling approach, data generation, and model evaluation are available in Weinstein et al (2019, 2020) and a brief summary is provided in S2. This model can be used directly to make predictions for new data or used as a foundation for retraining the model using labeled data from a new application.

**Figure 1.**
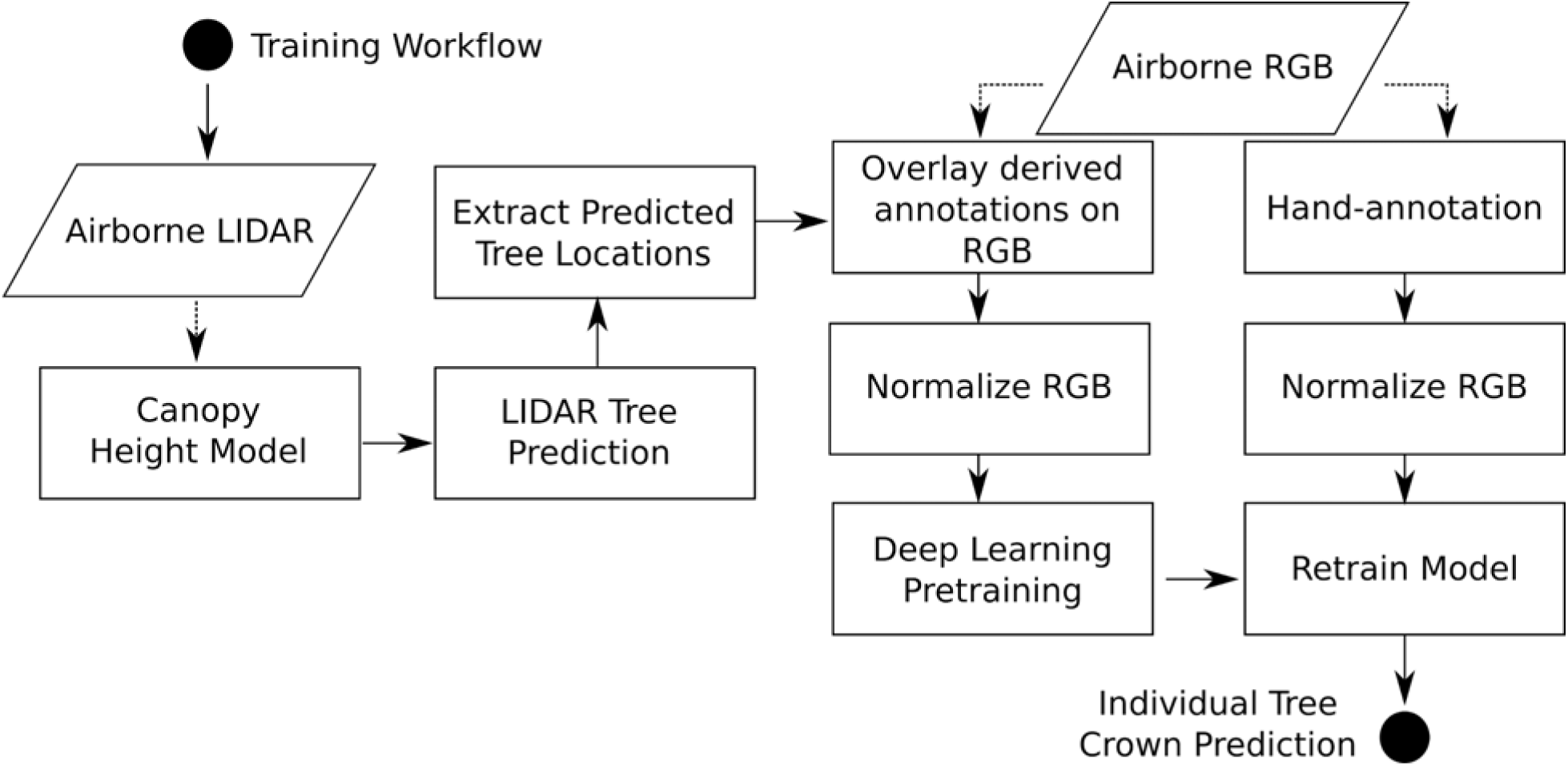
Prebuilt model training workflow. Redrawn from Weinstein et al. (2020). Parallelograms in the workflow indicate input data, rectangles indicate an algorithmic step, and circles indicate the start and end of the workflow. The two sub-flows on the right-side of the figure can run in parallel and outline the pre-training and fine-tuning stages of the overall model fitting process.

### Training

Tree crown delineation is a challenging task and a single model cannot realistically be expected to capture the tremendous taxonomic diversity at a global scale. This means that to perform optimal crown delineation for a particular forest requires training or fine-tuning using data from a local area. A key advantage of DeepForest’s neural network structure is that users can retrain the prebuilt model to learn new tree features and image backgrounds while leveraging information from the existing model weights based on data from a diverse set of forests. Fine-tuning neural networks starting from an initial model requires less training data to produce reasonable results (Shin et al. 2016). Known as “transfer learning”, this ability is important because training deep learning models from scratch often requires tens of thousands of labeled data points for ecological tasks (Weinstein 2018). In contrast, fine-tuning the prebuilt model with as few as 1000 hand labeled trees can provide significant improvement and be accomplished in approximately 8-10 hours (Weinstein et al. 2020).

The standard training process starts with generating local training data by hand-labeling trees in images by placing a bounding box around each visible tree (Figure 2). This can be done using either image labeling tools (e.g., RectLabel) or GIS software (e.g., ArcGIS, QGIS) and DeepForest includes helper functions to convert common formats (XML and shapefiles) into a csv format. Annotations can be made on images of any size, but training the model requires images with fixed standard dimensions. The prebuilt model was trained on square crops of length 400px (40m at 0.1m resolution), which provides a good balance between image size and providing the model landscape context for prediction. DeepForest includes a `preprocess.split_raster` function that creates a set of appropriately sized images for training using a sliding window approach. The size of these input windows are optimized for the 10cm data used in training the prebuilt model. The upper resolution limit for tree crown delineation is currently unknown, as well as the optimal size of the input windows when performing predictions at coarser scales.

**Figure 2.**
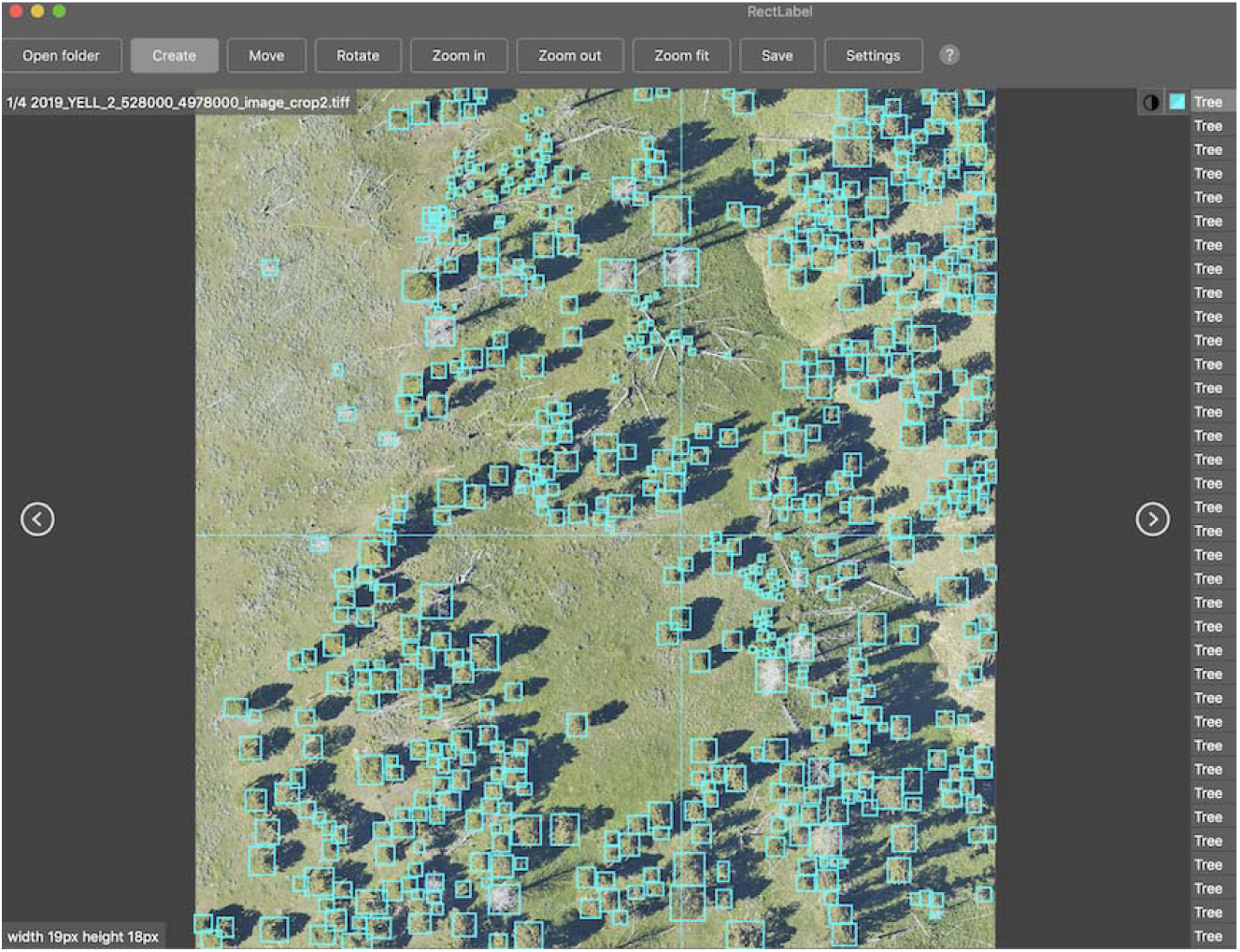
Screenshot of hand-annotated RGB image from NEON site YELL near Frog Rock, WY. For optimal training, all crowns in an image should be annotated.

Training can be performed by fine-tuning the prebuilt model or training only using the local training data (using `deepforest.train`). Training deep learning models requires a number of parameter choices such as batch size and number of epochs. For users less familiar with training deep learning models, DeepForest comes with a standard configuration file with reasonable defaults. While some parameter exploration will always be helpful, our aim is to make these innovations available even to novice users. Optional GPU support and model customization allow more experienced users to quickly develop and test larger and more complex models. Data augmentation to randomly crop and flip training images is also supported. This strategy is often useful to reduce overfitting when training on small datasets (Zoph et al. 2019) but has not been extensively tested for tree crown delineation. For additional recommendations for optimal model training see the online documentation (https://deepforest.readthedocs.io/) and Appendix S2.

### Evaluation

Deep neural networks have millions of parameters and can readily overfit, producing high scores on training data, while performing poorly on new images. This makes it essential to evaluate performance on held-out test data. To evaluate a set of annotations, users follow the same pattern as with training data: 1) Annotate one or more images of trees; 2) Cut the images into smaller windows for evaluation; and 3) Format annotations into a csv file using DeepForest’s utility functions. The `deepforest.evaluate_generator` method can then be used to evaluate the performance of the predictions for this test data using the mean average precision (mAP). mAP combines precision and recall into a single metric measuring the area under the precision-recall curve resulting in a score ranging from 0 to 1. In our experience, mAP scores above 0.5 are usable for scientific application, but the appropriate value depends on the particular research goal and application.

### Prediction

After a model has been trained and evaluated, it can be applied to a larger collection of images to estimate the locations of trees at larger scales. High resolution images covering wide geographic extents cannot fit into memory during prediction and would yield poor results due to the size and density of bounding boxes. DeepForest has a `deepforest.predict_tile` method for automating the process of splitting the tile into smaller overlapping windows, performing prediction on each of the windows, and then reassembling the resulting annotations. Each bounding box annotation is returned with its xmin, ymin, xmax, ymax coordinates, and predicted probability score (the probability that the bounding box represents a tree) ranging from 0-1, with higher values indicating greater confidence in the prediction. To reduce overcounting among overlapping tiles, DeepForest sorts predictions by confidence scores and removes lower scoring overlapping boxes (i.e., non-max suppression).

## Case Studies

### National Ecological Observatory Network

To evaluate model performance across a range of forest types, we used data from the National Ecological Observatory Network to predict crowns in sites across the United States. This dataset consists of 212 images containing 5852 trees from 22 sites that is part of an upcoming tree crown benchmark data package (Weinstein et al. 2020). Training and evaluation data are separated by at least 1 km when they occur at the same site. Evaluation data were created by viewing RGB images and manually delineating tree crown boxes for all visible trees. Annotations were cross-referenced with field collected positions of tree stems (from the NEON Vegetation Structure dataset; NEON ID: DP1.10098.001) within each plot when available. Following Weinstein et al. (2019, 2020), we used precision, defined as the fraction of predicted crowns match real trees, and recall, defined as the fraction of all evaluation trees that are correctly detected for evaluation. Following the standard evaluation for object detection in the computer vision literature (Ren et al. 2015), we considered predictions with Intersection over Union (IoU) scores of 0.5 as true positives. IoU, also known as the Jaccard Index, is the area of intersection between the prediction and evaluation crown, divided by the joint area of the combined prediction and evaluation crowns. We assessed the performance of the prebuilt model at all 22 NEON sites and also compared the performance to a previous version of this model (Weinstein et al. 2020) that was only trained on data from 4 NEON sites.

Across all sites the average recall per image for the prebuilt model was 72% and the precision was 64%. Model performance varies across NEON sites, but most sites have both precision and recall values greater than 50% (Figure 3). The model performs similarly regardless of whether there was hand-annotated training data from the same site (Figure 3). Visual assessment of predictions across forest types reveals good overall correspondence between predicted bounding boxes and observations, with most errors resulting from insufficient overlap between observed and predicted tree crowns, rather than the model missing a tree entirely (Figure 4). The prebuilt model used by DeepForest was fit to data from 22 NEON sites and outperforms the previous 4 site model (Weinstein et al. 2020) at 19 of 22 sites for recall and 16 of 22 sites for precision, demonstrating that increasing the diversity and amount of training data has improved the performance of the model. These results demonstrate that the prebuilt model can make reasonable predictions in forests ranging from deciduous forests of the Northeast, to southern pinelands, to coniferous forests of the mountain west.

**Figure 3.**
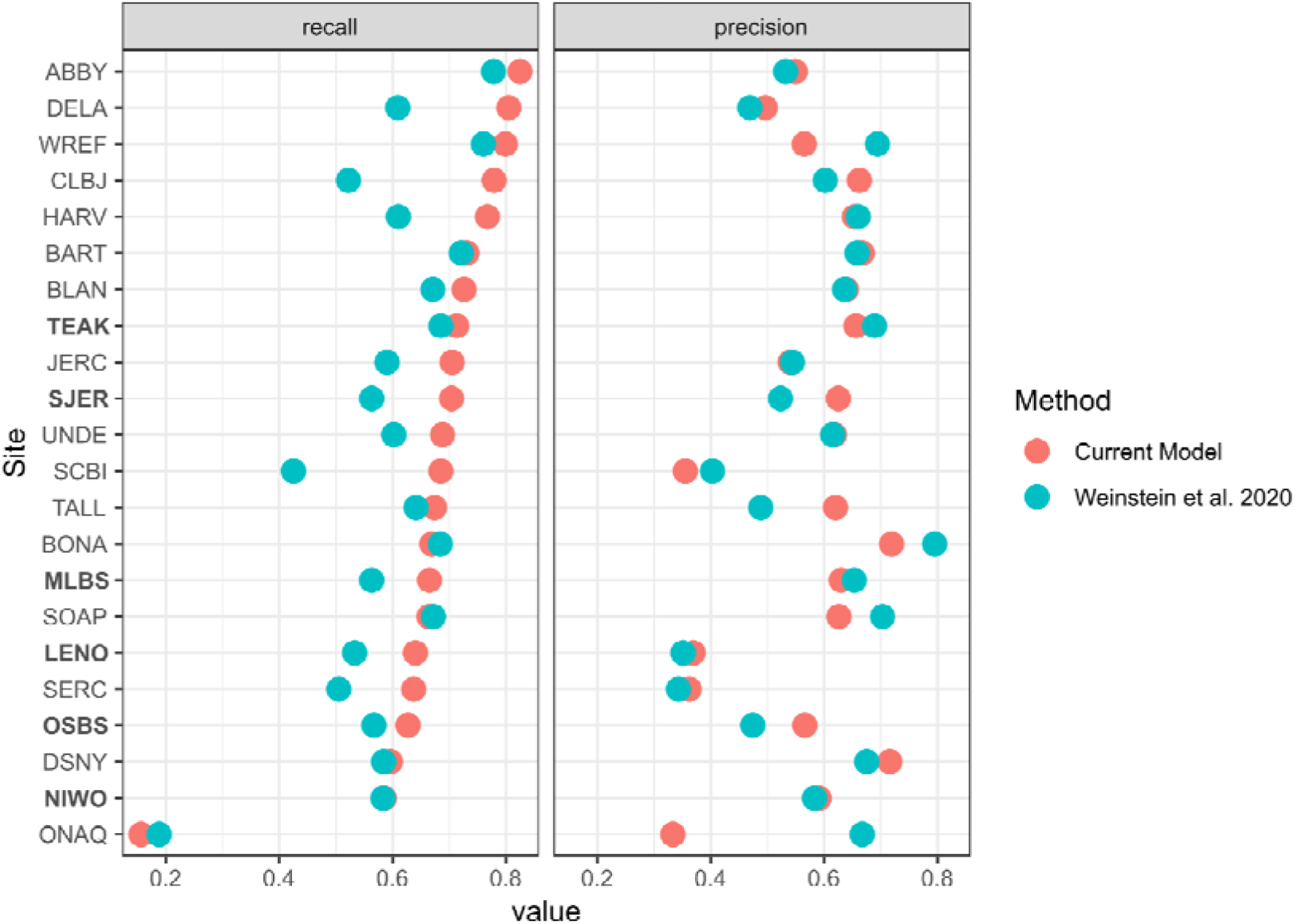
Precision and recall scores for hand-labeled evaluation images from the National Ecological Observatory Network (current prebuilt model in red, Weinstein et al. 2020 in blue). Sites in bold had hand-labeled data included in training the current prebuilt model. See S1 for site abbreviations.

**Figure 4.**
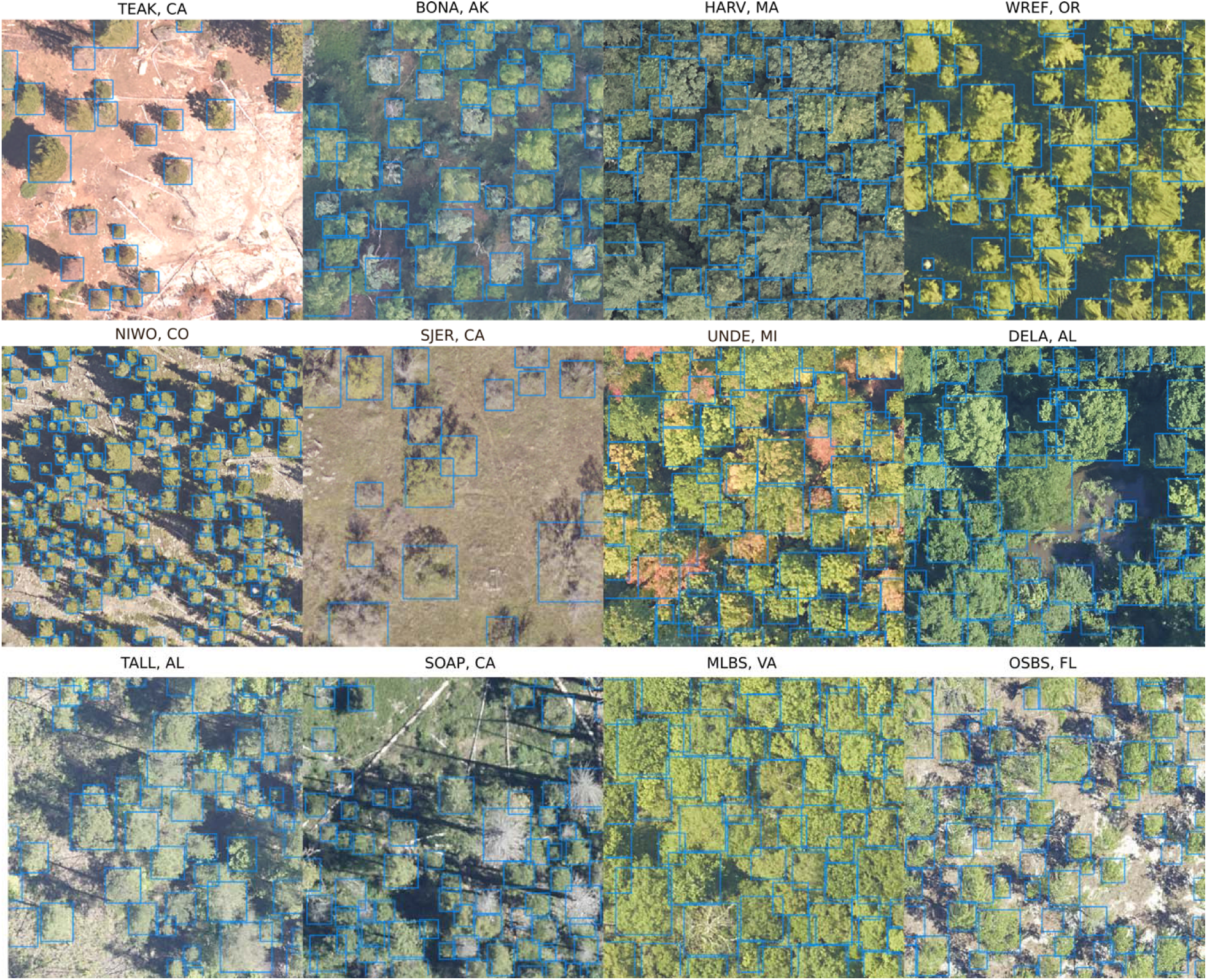
Panel of tree predictions from a broad range of evaluation images in the National Ecological Observatory Network with predicted tree crown boxes are in blue. Each image is labeled with the NEON site abbreviation and state. See S1 for site abbreviations.

The site with the worst performance is Onaqui, Utah (ONAQ), which is a desert scrub site with a different vegetation structure from any of the training data. The site is almost treeless and includes trees with short and gnarled stature. This highlights the importance of using local training data to reduce uncertainty when working with data that is not well represented in the training data for the prebuilt model. In these contexts, the value of the prebuilt model is that it reduces the needed training sizes when applied to new conditions. This has the potential to support training with small amounts of data for applications to a wide array of questions surrounding tree health and ecology. For example, training a model specific to bare trees could allow studies of broad-scale pest outbreaks or timing of deciduous phenology. Initial tests at the Soaproot Saddle, CA site (‘SOAP’ in Figure 4, 3^rd^ row) show the prebuilt model can detect standing dead trees when visible. Adding additional training data could allow broad scale analysis of tree health when comparing images across time.

### French Guiana Tropical Forest

The DeepForest prebuilt model was trained on data from the United States that was collected using fixed-winged aircraft at 10cm resolution and provided as 1km2 orthomosaics. Therefore, two key questions are: 1) Does this model generalize to images collected in new locations or using different acquisition hardware; and 2) how useful are the (re)training features of the software for improving performance in novel contexts? It is also important to understand how the DeepForest RGB model compares to LiDAR-based models from recently published work.

To address these questions, we used data from a recently published competition comparing LIDAR tree segmentation algorithms using remote sensing from French Guiana (Aubry-Kentz et al. 2019). In the original competition, each team was sent unlabeled data to predict and the evaluation data was kept private. This process was repeated for this paper, with the third author (Aubry-Kientz) running evaluation scores for the DeepForest predictions made by the corresponding author (Weinstein). Predictions were run on a Mac laptop with a 3.1 GHz Intel Core i5 processor. Predictions from each algorithm were compared to hand-delineated evaluation crowns based on field observation and manual comparison with RGB and LiDAR data. Validation crowns were delineated as polygons, rather than the rectangular bounding boxes generated by DeepForest. This case study also provides information on whether DeepForest’s approach of predicting rectangular bounding boxes leads to lower prediction accuracy than methods producing polygons. Algorithm recall was scored based on the proportion of labeled trees predicted with IoU scores of greater than 0.5. Precision was not calculated because not all crowns in the test imagery were delineated (see Figure 5).

**Figure 5.**
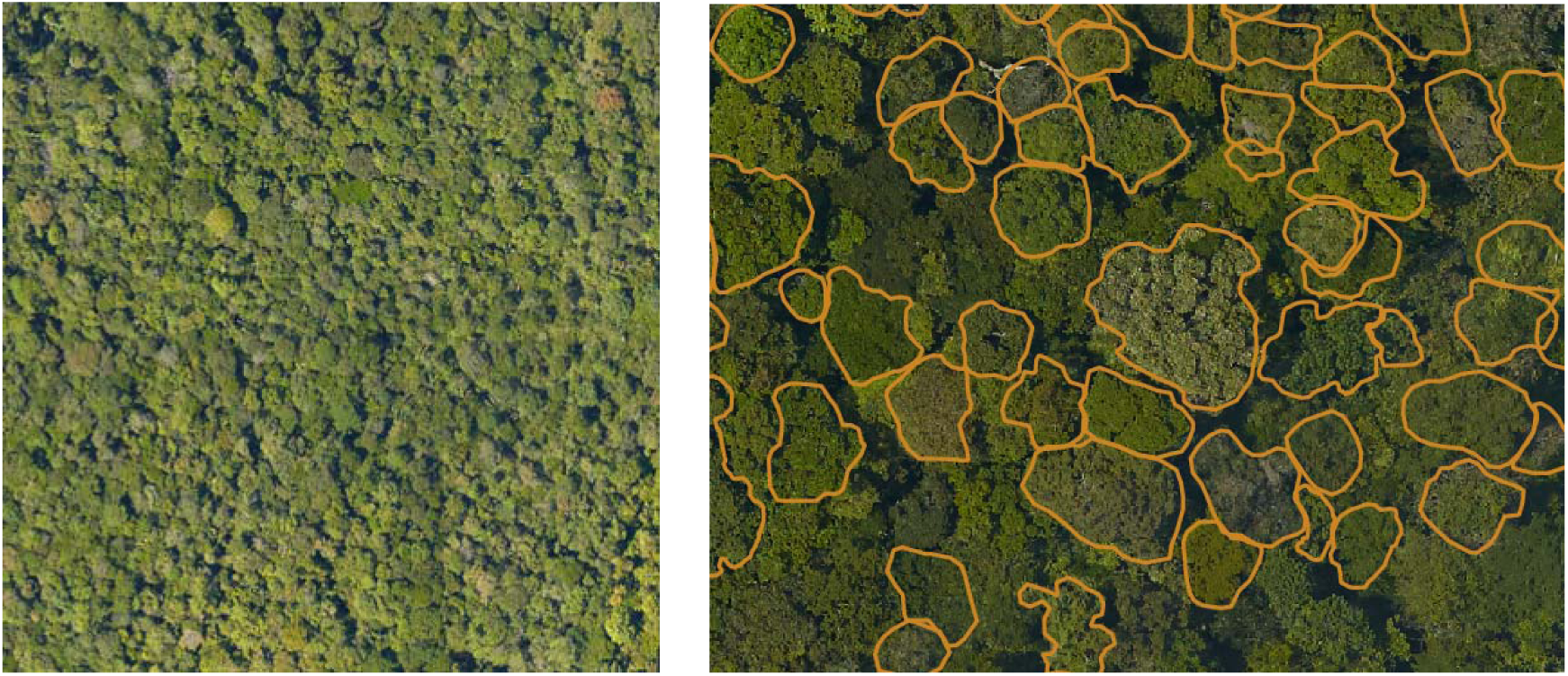
RGB images collected over a tropical forest in French Guiana and example of manually segmented crowns used to evaluate the segmentation.

We used DeepForest to detect tree crowns using three approaches 1) the prebuilt DeepForest model with no local training; 2) a model fit solely to 5018 local hand-annotated crowns (annotated by BW using only the RGB data on tiles separate from the evaluation data) and 3) the prebuilt model fine-tuned using the local annotations. RGB tiles were divided into 800px windows for model training and evaluation. The default patch size of 400px was increased to 800px to minimize the edge effect of overlapping crowns. Models 2 and 3 were trained for 7 epochs with a runtime of approximately 11 minutes/CPU on a laptop, which demonstrates that while advanced GPU hardware is convenient for training large datasets, fine-tuning and training on small datasets can be done locally on CPU.

The prebuilt model performed well on this novel data with a recall of 0.64, close to the 0.71 recall for the best performing LIDAR based algorithm from Aubry-Kientz et al. (2019). Training only on the 5018 local annotations resulted in a poorer recall of 0.35Retraining the prebuilt model with the local annotations produced the best results with a recall of 0.78, slightly better than the highest performing LiDAR algorithm from Aubry-Kientz et al. (2019). This analysis is not sufficient to draw general conclusions about RGB versus LiDAR-based methods, but these results do suggest that DeepForest is competitive with state-of-the-art LiDAR-based approaches. Overall, the case study demonstrates the utility of DeepForest both using the prebuilt model and using local retraining to improve crown delineation based on local conditions.

### Portland Street Trees

DeepForest’s use of widely available RGB data provides the potential for it to be used across very large spatial extents. Scaling up is challenging because algorithms need to handle large ranges of habitat types and because the resolution of the data available over large areas is typically coarser. To explore how DeepForest performs using coarser resolution data in unique habitats, we applied both the prebuilt model and a retrained model to crown delineation of street trees in an urban environment. The locations of urban trees are important for ecological, sociological and public infrastructure applications. In addition, the urban environment is very different from the natural environments on which the prebuilt model was trained. The image data from the Oregon Statewide Imagery Program is also coarser at 0.3m spatial resolution (1ft), a resolution that is widely available as part of the National Agriculture Imagery Program (NAIP - https://www.fsa.usda.gov/programs-and-services/aerial-photography/imagery-programs/naip-imagery/).

We used imagery from the Portland metro area that overlapped with the Portland Street Trees dataset (http://gis-pdx.opendata.arcgis.com/datasets/street-trees). The street trees dataset contains geospatial information for the majority of trees accessible from public roads in the metro area. Not all trees in an image are labeled, since many trees occur on private property and are not mapped. We divided the RGB imagery into geographically distinct training and test datasets and used the street trees dataset to guide hand-annotation of a small number of tree crowns (n=1033). Annotation by hand took approximately three hours and covered a small geographic area of mixed urban development, empty lots and ballfields (Figure 6). The street trees data was collected prior to the RGB images and was cleaned to remove trees that had been cut down or were obvious errors (e.g. trees located in the middle of buildings). To evaluate the street tree case study, we used field collected location of the tree stems to measure tree recall and the rate of undersegmentation. Recall was defined as the proportion of street tree locations that were contained within a predicted tree bounding box. Undersegmentation rate was defined as the proportion of predicted boxes that matched more than one street tree. Minimizing undersegmentation is challenging because trees growing close together can appear to be a single tree from above and is therefore best evaluated against ground collected data.

**Figure 6.**
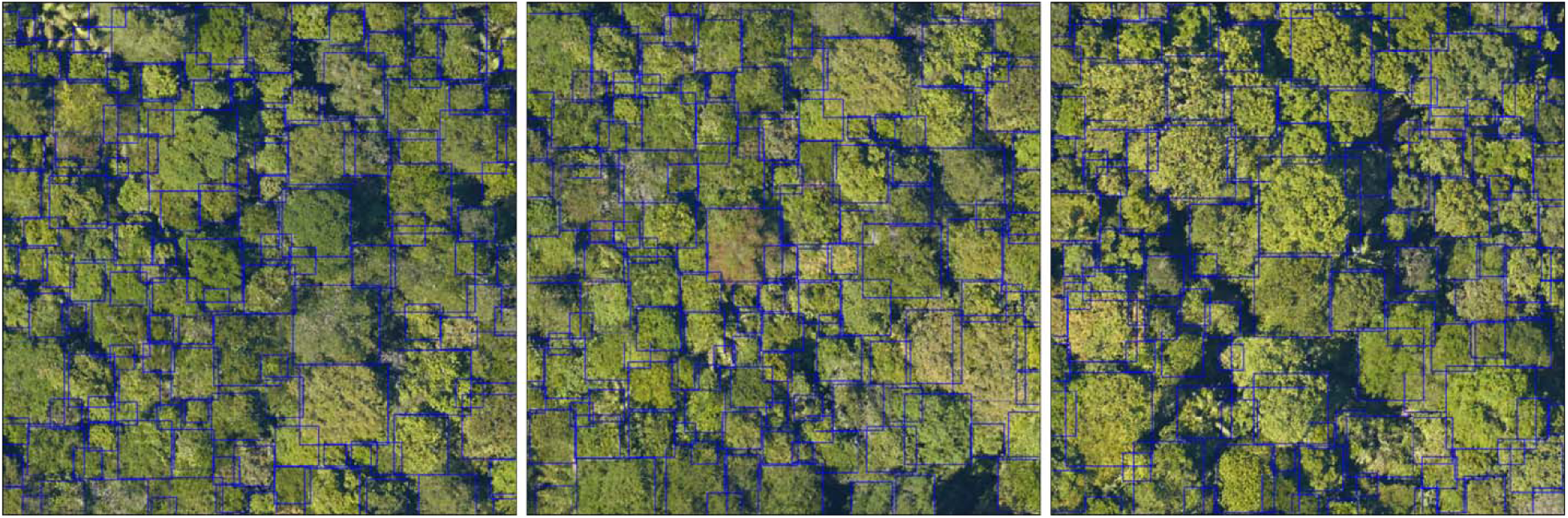
Predictions made on a tropical forest in French Guiana using the prebuilt model retrained with local annotations. Each individual tree is labeled with a blue bounding box.

We found that evaluating and retraining on data with coarser resolution than the prebuilt model required careful choosing of the size of the focal view. The prebuilt model was originally trained on a 40m focal view (400px windows with 0.1m data). Data exploration on the coarser data source showed that larger focal views of 60-120m performed better than maintaining the original 40m view, and 60m was chosen for this analysis. In general, we expect that the focal view size should increase with coarser resolution data, but this remains an area of further exploration.

As with the tropical forest case study, we found that the prebuilt model performed reasonably well (recall = 0.55; undersegmentation = 0.25) and retraining with a small amount of local training data significantly improved algorithm performance with an increase in recall and decrease in undersegmentation (recall = 0.72; undersegmentation = 0.17; Figure 7). Visual inspection shows that many of the errors in using the retrained model are for small trees difficult to resolve in the imagery, or tree types not present in the limited training data (e.g. ornamental trees with a deep purple hue).

**Figure 7.**
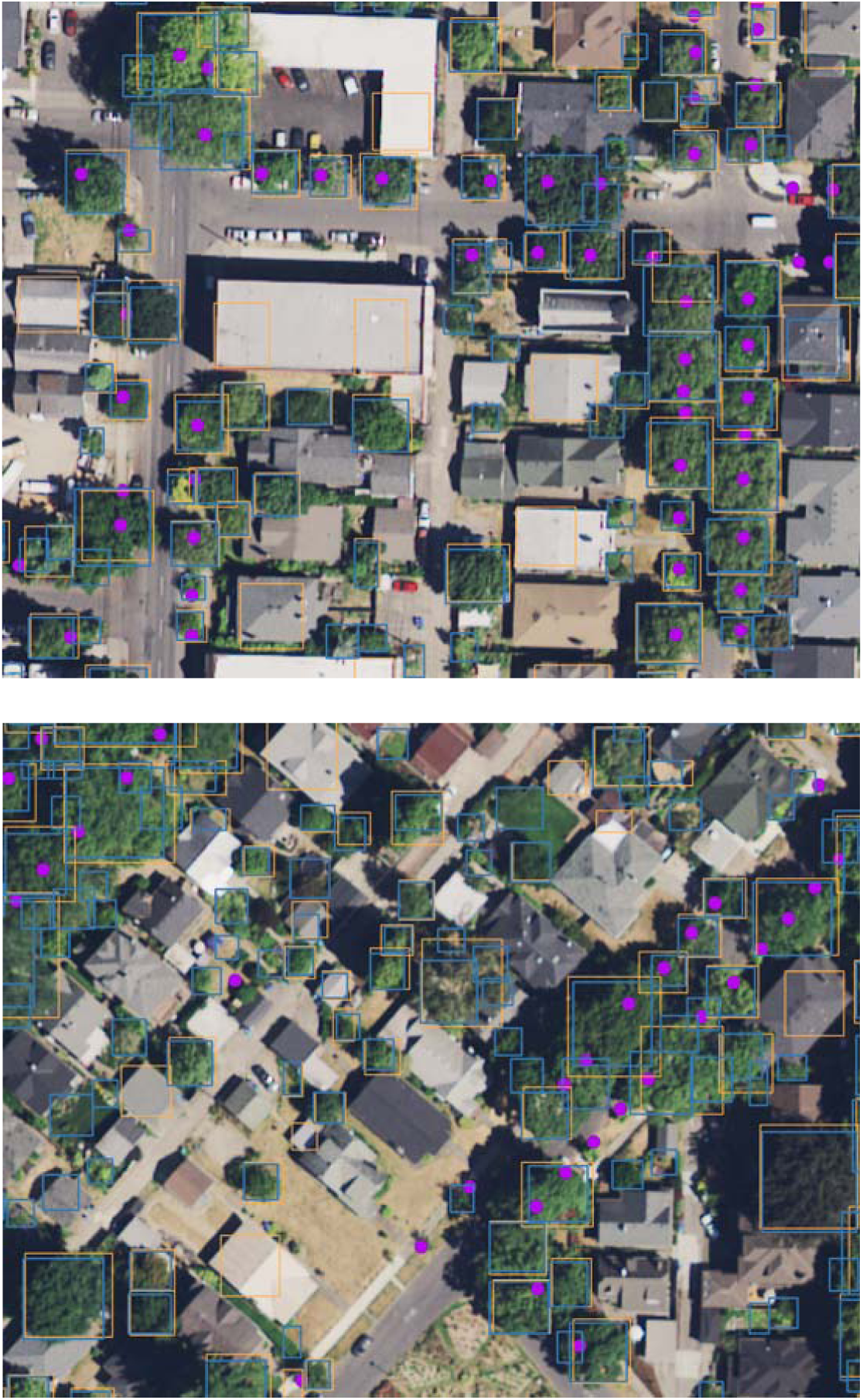
Predictions for the Portland street tree case study. Bounding box predictions from the prebuilt model are in orange. Bounding box predictions from the retrained model using local data are in blue. Street tree locations are marked in purple.

## Conclusion

DeepForest provides an open source software package for: 1) delineating tree crowns in RGB imagery, 2) evaluating the performance of that crown delineation using hand labeled evaluation data, and 3) training new models and fine-tuning of the included prebuilt model to support prediction tailored to specific forest types. The inclusion of a prebuilt model allows users to benefit from the strengths of deep learning without needing to deal with many of the challenges. Given the enormous diversity of tree appearance at a global scale, defining a single unified model for tree crown delineation is challenging. To address this, DeepForest provides an explicit retraining method to improve performance for specific use cases. This allows the user to decide what level of accuracy is required for the target question, and then annotate local data and retrain the model to produce predictions with sufficient accuracy for their use case. We recommend defining a clear evaluation dataset, setting a threshold for desired performance before training, and using evaluation data that is geographically separate from the training data to ensure that the prediction threshold holds outside of the training region.

The minimal spatial resolution for accurate tree prediction using this software remains unknown and may ultimately relate to the desired ecological or management question. Analysis of the NEON data show that individual tree segmentation is achievable at 10cm. The Portland Street trees example shows that 30 cm data (which is publicly available for many states and counties) provides reasonable delineations. However, the accuracy will not be as high as with higher resolution data, and further analysis at this resolution is necessary. One meter resolution imagery is increasingly available at near continental scales (e.g., NAIP 1m imagery which provides nearly complete coverage of the United States; https://www.fsa.usda.gov/programs-and-services/aerial-photography/imagery-programs/naip-imagery/). It is unlikely that these data will be effective at distinguishing small individual trees, but it may be useful in identifying large trees or clusters of trees in sparse landscapes.

To support the broad application of predictions from DeepForest, these predictions can be easily exported for use in further analysis and combination with other sensor products for forest research. Individual tree crown delineation is often the first step in key remote sensing analyses of forested landscapes, including biomass estimation (Kamoske et al. 2019), species classification (Maschler et al. 2018), and leaf-trait analysis (Marconi et al. 2019). DeepForest both ingests and outputs crowns in an easily accessible, standardized annotation format, and will facilitate further improvements in the prebuilt model based on community contributions.

## Data Availability

DeepForest source code is available on GitHub (https://github.com/weecology/DeepForest) and archived on Zenodo (https://doi.org/10.5281/zenodo.2538143). The code for the case studies is available in a separate repo (https://github.com/weecology/DeepForest_demos). The in-development version of the NEONTreeEvaluation benchmark is available online (https://github.com/weecology/NeonTreeEvaluation) and will continue to be updated as more images are annotated. The Oregon RGB imagery was provided by the Oregon Statewide Imagery Program 2018: 1 foot orthophotography of western Oregon: State of Oregon data release, https://www.oregon.gov/geo/Pages/imagery_data.aspx

## Author Contributions

BW, SM, EW conceived of the project, designed the package and wrote the manuscript. MAK, GV, and HS performed analysis, provided package improvements and edited the manuscript.

## Acknowledgments

We thank Brady Callahan for his assistance in accessing Oregon imagery data and the Portland Metro for their work collecting the Portland Street Trees dataset. This research was supported by the Gordon and Betty Moore Foundation’s Data-Driven Discovery Initiative through grant GBMF4563 to E.P. White and by the National Science Foundation through grant 1926542 to E.P. White, S.A. Bohlman, A. Zare, D.Z. Wang, and A. Singh. We thank Alina Zare, Aditya Singh, and Dylan Sinnickson for feedback during model and software development and comments on the manuscript. The authors declare no conflicts of interest.

---

**Supplementary 1.** NEON Site Abbreviations

| *Site Name* | *Site ID* | *State* | *Latitude* | *Longitude* |
| --- | --- | --- | --- | --- |
| *Onaqui* | ONAQ | UT | 40.17759 | -112.45244 |
| *Santa Rita Experimental Range* | SRER | AZ | 31.91068 | -110.83549 |
| *Niwot Ridge Mountain Research Station* | NIWO | CO | 40.05425 | -105.58237 |
| *Yellowstone Northern Range (Frog Rock)* | YELL | WY | 44.95348 | -110.53914 |
| *Wind River Experimental Forest* | WREF | WA | 45.82049 | -121.95191 |
| *Abby Road* | ABBY | WA | 45.76243 | -122.33033 |
| *Lower Teakettle* | TEAK | CA | 37.00583 | -119.00602 |
| *Caribou-Poker Creeks Research Watershed* | BONA | AK | 65.15401 | -147.50258 |
| *Soaproot Saddle* | SOAP | CA | 37.03337 | -119.26219 |
| *San Joaquin Experimental Range* | SJER | CA | 37.10878 | -119.73228 |
| *Lenoir Landing* | LENO | AL | 31.85388 | -88.16122 |
| *UNDERC* | UNDE | MI | 46.23388 | -89.53725 |
| *Ordway-Swisher Biological Station* | OSBS | FL | 29.68927 | -81.99343 |
| *Disney Wilderness Preserve* | DSNY | FL | 28.12504 | -81.4362 |
| *Jones Ecological Research Center* | JERC | GA | 31.19484 | -84.46861 |
| *Blandy Experimental Farm* | BLAN | VA | 39.06026 | -78.07164 |
| *Smithsonian Environmental Research Center* | SERC | MD | 38.89008 | -76.56001 |
| *Smithsonian Conservation Biology Institute* | SCBI | VA | 38.89292 | -78.1395 |
| *Bartlett Experimental Forest* | BART | NH | 44.06388 | -71.28731 |
| *Konza Prairie Biological Station* | KONZ | KS | 39.10077 | -96.56309 |
| *Talladega National Forest* | TALL | AL | 32.95046 | -87.39327 |
| *Dead Lake* | DELA | AL | 32.54172 | -87.80389 |
| *Mountain Lake Biological Station* | MLBS | VA | 37.37828 | -80.52484 |
| *The University of Kansas Field Station* | UKFS | KS | 39.04043 | -95.19215 |
| *Harvard Forest* | HARV | MA | 42.5369 | -72.17266 |

S2: Technical background to DeepForest and advice on model development

The goal of this paper is to provide a robust open-source implementation for RGB tree crown delineation. For detailed information on data generation and model testing see Weinstein et al. (2019, 2020). Here we provide an overview with recommendations for parameters that could affect model performance. For more discussion with code examples see https://deepforest.readthedocs.io/.

DeepForest is a deep learning object detection model. Deep learning uses a series of hierarchical filters to connect low-level image features, such as colors and shape, to high-level concepts such as crown contours. Deepforest uses the keras-retinanet implementation (Gaiser et al 2017) of the Retinanet one-stage object detector with a Resnet-50 classification backbone (Ren et al. 2015). This classification backbone was pretrained on the ImageNet dataset (He et al., 2016). This pretraining is useful for reducing training time and minimize overfitting to smaller ecological datasets. Given an input image, the model predicts both the location of bounding boxes and object classes simultaneously. In the DeepForest implementation, the prebuilt model has only 1 “Tree” class. All instances of the target class are predicted, such that if there are 40 trees in the image, an ideal model will predict 40 bounding boxes. Deep learning models have millions of parameters and take significant computational resources. To reduce memory consumption, we cut each prediction image into 40m by 40m windows with an overlap of 5%. Using a pool of unsupervised LiDAR-based tree predictions generated using Silva et al (2016), we pretrained the network with a batch size of 20 on 2 Tesla K80 GPU for 5 epochs. To align these unsupervised classifications with the ImageNet pretraining weights, we normalized the RGB channels by subtracting the ImageNet mean from each channel. We then retrained the network using the hand-annotated data for 40 epochs. Data augmentation of random flips and translations was tested and found to have little effect on the final score.

## Key parameter choices

Given the diversity of forests, data resolutions and image backgrounds, it is unlikely that a universal tree crown method exists without some parameter tuning. The prebuilt model does a good job in many situations, but to get the most of of DeepForest, some parameter exploration will be needed. Using the prebuilt model, users should consider:

### Patch Size

The prebuilt model was built on 40m crops with 0.1m imagery (400^2 pixels). The model can generalize to new resolutions, (see street trees example), but may need to vary the image patch size. For example, on the 30cm NAIP data, we found that increasing patch size from 40 to 60 meters per image improved model performance. While an exact equation between image resolution and optimal patch size is unknown, we anticipate that coarser resolution data require larger patches to gather more context around visible tree crowns.

### IoU suppression threshold

Object detection models have no inherent logic about overlapping bounding boxes. For tree crown detection, we expect trees in dense forests to have some overlap, but not be completely intertwined. We therefore apply a postprocessing filter called ‘non-max-suppression’, which is a common approach in the broader computer vision literature. This routine starts with the boxes with the highest confidence scores and removes any overlapping boxes greater than the intersection-over-union threshold (IoU). Intersection-over-union is the most common object detection metric, defined as the area of intersection between two boxes divided by the area of union. If users find that there is too much overlap among boxes, increasing the IoU will return fewer boxes. To increase the number of overlapping boxes, reduce the IoU threshold.

## Training New Models

Ultimately, the best performance will come from training local data. Even a small amount of local training data can be beneficial. When training images, it is important to label all visible trees, not just the trees that the user has high confidence in. Skipping trees in an image will teach the model to skip trees during prediction. During annotation, we highly suggest users spend the time to answer 2 questions: 1) What kind of data am I trying to predict? Capturing the variability and the broad range of tree taxonomy and presentation will make development go more smoothly. 2) What kind of accuracy do I need to answer my question? It is natural to want the best model possible, but one can waste a time trying to eek out another 5% of recall without understanding whether that increase in performance will improve our understanding of a given ecological or natural resource question.

### Batch size

Neural networks are often trained in batches of images, since the entire dataset is often too large to read into memory at once. The size of these batches affects both the speed of training (larger batches train faster) and the stability of training (larger batches lead to more consistent results). The default batch size of 1 is chosen because it is not possible to anticipate the available memory and should be increased whenever possible. Typically, batch sizes are evenly divisible by the size of the entire dataset.

### Epochs

An ‘epoch’ is one iteration of training on all images. The more times the model sees the training data, the higher the training accuracy. However, deep learning models have millions of parameters and can easily overfit. For small datasets (< 10,000 annotations), 5-10 epochs should be sufficient. If training for longer, make sure to carefully evaluate validation accuracy on out-of-sample data.

## Notes

https://deepforest.readthedocs.io/

### Competing Interest Statement

The authors have declared no competing interest.

